# Tick-Borne bacterial and protozoan animal pathogens shape the native microbiome within *Hyalomma anatolicum anatolicum* and *Rhipicephalus microplus* tick vectors

**DOI:** 10.1101/2020.01.20.912949

**Authors:** Abdulsalam Adegoke, Deepak Kumar, Muhammad Imran Rashid, Aneela Zameer Durrani, Muhammad Sohail Sajid, Shahid Karim

## Abstract

**Background:** Ticks vector a variety of bacterial, viral, and protozoan pathogens of public and animal health significance. Ticks also harbor a diverse community of microbes linked with their biological processes like hematophagy and hence vector competence. The interactions between bacterial and/or protozoan pathogens and their tick vector microbiome are yet to be investigated. In lieu of this, this study was designed to define the microbial composition of uninfected and infected *Hyalomma (H.) anatolicum anatolicum* and *Rhipicephalus (R.) microplus* tick species.

**Methodology/Principal findings:** A total of 320 *H. anatolicum* and *R. microplus* were screened for the presence of the protozoan (*Theileria* sp.), and bacterial (*Anaplasma marginale*) pathogens by PCR. Subsequently, the microbiome of uninfected and infected individual *H. anatolicum* and *R. microplus* were analyzed. The highly conserved V1-V3 region of the 16S rRNA gene was sequenced using the MiSeq Illumina platform. The microbiome of female *H. anatolicum anatolicum* ticks was dominated by the endosymbiont *Candidatus Midichloria mitochondrii* (CMM) and *Francisella*-like endosymbiont (FLE) which were not affected by pathogen infection. *Ehrlichia* species was detected in *A. marginale*-infected male *H. anatolicum anatolicum* (6.2%) as opposed to the *Theileria* sp*.-*infected female *H. anatolicum anatolicum*. *Coxiella* sp. was also detected in uninfected (2.96%) and *A. marginale*-infected (4.25%), but not in *Theileria* sp.-infected *R. microplus* ticks. Analysis of the eukaryote composition in the respectively ticks also revealed the presence of operational taxonomic units (OTUs) belonging to *Plasmodium (P.) falciparum* in *Theileria* sp.-infected *H. a. anatolicum* and *R. microplus* ticks, while *Hepatozoon americanum* detected from *Theileria* sp.-infected and uninfected *H. a. anatolicum*.

**Conclusion and Significance:** This study establishes the extent of the diversity of microbial community of two important tick species from Pakistan and also revealed the presence of *Theileria* and *A. marginale* and additional pathogenic bacteria that could be of public health significance. We hypothesized that infection with either a protozoan or bacterial pathogen will alter the microbial composition within these tick species. Interestingly, we reported the detection of the malarial parasite (*P. falciparum*) from ticks infected with the protozoan pathogen (*Theileria* sp.). Further validation experiments are required on endosymbionts and pathogens of ticks to investigate how they could be important in the epidemiology of human and animal pathogens.

## Introduction

Ticks are known obligate, blood feeding ectoparasites of vertebrate animals that depend on the host’s blood to carry out nutritive, reproductive and other physiologic functions. In addition to causing significant blood loss, they are also known to transmit infectious pathogens such as viruses, bacteria, protozoa, and fungi to their animal or human hosts during the feeding process. Recent studies have demonstrated ticks harbor microbial communities, members of which undergoes mutualistic and commensal relationships with their tick hosts, most of which have been commonly overlooked or considered as potential tick-borne pathogens (1–5). Therefore, it is posited that a constant wave of interaction will continually ensue between tick-borne pathogens (TBPs) and bacterial endosymbionts. TBPs and bacterial endosymbionts could directly compete for nutrient or niche within the tick hosts. Indirect inhibition or elimination could also occur through excretory molecules inhibiting growth of a competitor (6). With this knowledge comes the understanding that the microbial communities including TBPs must continuously co-evolve to maintain equilibrium within the tick vectors.

The most reported tick-borne disease in Pakistan includes Theileriosis, Babesiosis and Anaplasmosis caused by *Theileria (T.) annulata, Babesia (B.) bigemina, B. divergens, Anaplasma (A.) marginale and A. centrale*, which have all been found to affect water buffaloes and cattle (7–8, 54). The maintenance of these diseases has been made possible due to the availability and endemicity of suitable tick vectors belonging to the Ixodid group (9, 55). These include *H anatolicum, H. dromedarii, H. marginatum* as possible vectors of *T. annulata*; *R. microplus* as possible vectors for *B. bigemina, B. bovis, B. orientalis, B. divergens,* and *A. marginale* while *Hyalomma* sp. as possible vector for *A. marginale* and *A. centrale* (9).

While recent studies have been centered on TBPs, little attention has been paid to the entire microbial community, with only one such study determining the microbiome and pathogen diversity in ticks from Pakistan (10). These non-pathogenic microorganisms were either overlooked or considered to be potential TBPs (11). However, these non-pathogenic bacterial communities could confer multiple beneficial or detrimental effects on the tick host, interfere with the basic reproductive and fitness functions, and also alter the dynamics of TBP transmission (11).

A number of studies on the epidemiology and distribution of tick-borne pathogens have been published in Pakistan (7, 9–10, 17–18), there is a dearth of information regarding the microbiome of these ticks and how they interact with these pathogens within the same tick host. Here, we report the microbial communities residing within the uninfected, *Theileria* species and *Anaplasma marginale*-infected ticks.

## Results

### Pathogen prevalence

A total of 320 hard ticks pulled from cattle, sheep, and goat comprising of 198 *H. anatolicum anatolicum* and 122 *R. microplus*. All ticks were screened for the presence of *Theileria* sp. and *A. marginale* by PCR amplification using primers specific for the 18S rRNA and 16S rRNA gene respectively. Of the 320 samples, 23 (7%) and 90 (28%) were *A. marginale* and *Theileria* sp. positive respectively.

*A. marginale* had a prevalence of 2.6% (n=5) and 16% (n=23) in *H. anatolicum anatolicum* and *R. microplus* ticks, while the prevalence of *Theileria* was 36% (n=72) and 14% (n=18) in *H. anatolicum anatolicum* and *R. microplus* ticks (Table 1). The PCR amplicon was sequenced, and the nucleotide homology was assessed by searching the non-redundant nucleotide collection at GenBank (Supplementary data; Table1).

**Table 1:** Prevalence of *Anaplasma marginale* and *Theileria* species in *Hyalomma anatolicum anatolicum* and *Rhipicephalus microplus*

| Tick species | Numbers | *Anaplasma marginale* infected | *Theileria* species infected |
| --- | --- | --- | --- |
| *H. anatolicum anatolicum* | 198 | 5 (2.6%) | 72 (36%) |
| *R. microplus* | 122 | 20 (16%) | 18 (14%) |
| Total | 320 | 23 (7%) | 90 (28%) |

**Table 2:**
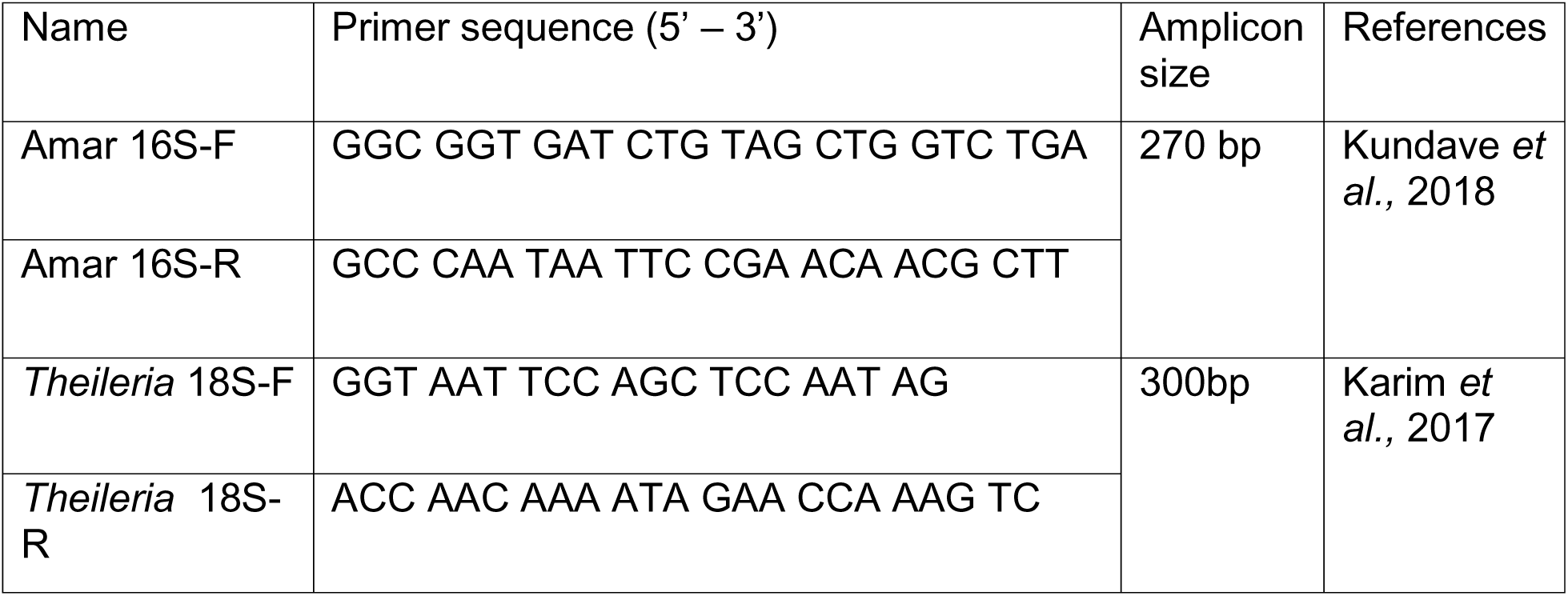
List of primers used in this study

### Microbiome composition

Analysis of the demultiplexed paired-end-reads generated 2902862 reads which ranged from 17063 to 123731 with an average of 65609 reads. Sequences from samples of *R. microplus* ticks generated the most reads of 1594076 while 1308786 reads belonged to sequences from *H. anatolicum anatolicum*. Taxonomic classification using the SILVA reference base identified 472 OTUs generated from *Rhipicephalus* ticks belonging to 10 phyla, 17 classes, 146 genera, and 137 species. *H. anatolicum anatolicum* had a total of 1314 OTUs representing 15 phyla, 33 classes, 196 genera, and 174 species.

#### Hyalomma anatolicum anatolicum

The phylum Proteobacteria was the most abundant in both *Theileria sp.-*infected and uninfected ticks (∼70%). The phylum Firmicutes had much more abundance within the *Theileria*-infected ticks representing about 20% of the total OTUs in the infected ticks. Other OTUs found to be present in relative amounts include phyla Actinobacteria, Tenericutes, and Bacteroidetes. OTUs with relative abundance below 1% were grouped into others. Similarly, the microbiome of *A. marginale*-infected male *H. anatolicum anatolicum* was dominated by the phylum Proteobacteria (∼48%) followed by the phyla Actinobacteria (24.5%), Firmicutes (18.9%), Bacteroidetes (4.7%) and Cyanobacteria (1.4%). In contrast, the representative phyla in uninfected male *H. anatolicum anatolicum* were evenly distributed among the phyla Proteobacteria, Actinobacteria and Firmicutes, while phyla Bacteroidetes and Cyanobacteria had the least abundance (Fig. 1A).

**Figure 1:**
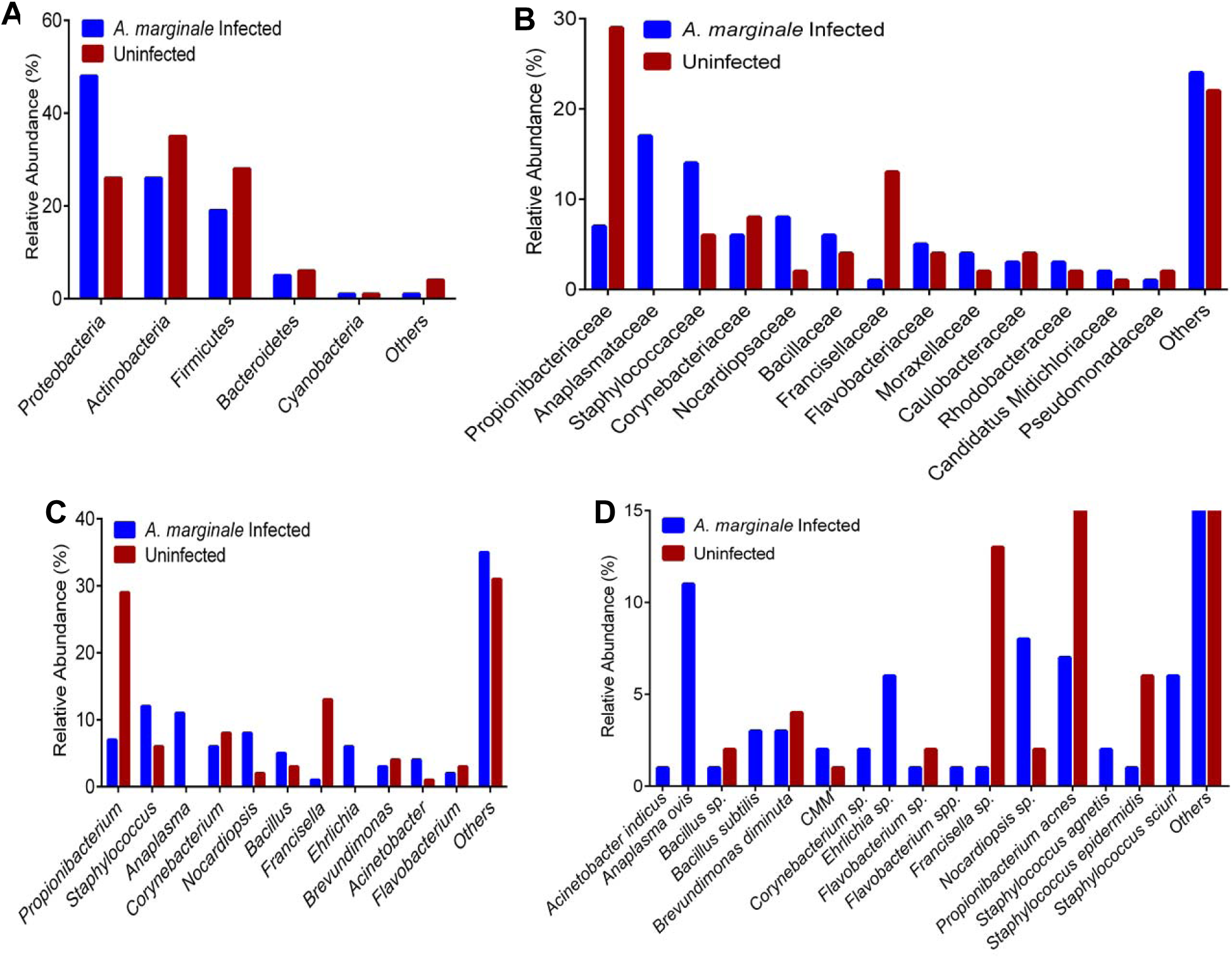
Relative abundances of taxa in *A. marginale* infected and uninfected male *H. anatolicum anatolicum* ticks at the A), phylum, B), family, C), genus and D), species level. Taxa with less than 1% abundance were grouped as others. 16S analysis revealed exclusive presence of *A. ovis*, *Staph. sciuri* and *Ehrlichia* species in PCR positive *A. marginale* ticks, indicating coinfection.

Uninfected male *Hyalomma* ticks had an abundance of *Propionibacterium* (29%), followed by *Francisella* (12.7%), *Corynebacterium* (8.2%) and *Staphylococcus* (6.11%) (Fig. 1C). With *A. marginale* infection, there was an observed increase in the number of representative genera within the female *Hyalomma* ticks with an even distribution a *Propionibacterium* (7.09%), *Staphylococcus* (12%), *Anaplasma* (10%), and the genus *Ehrlichia* (6.2%). The microbiome of both *Theileria-*infected and uninfected female *Hyalomma* ticks were dominated by the genus *Candidatus Midichloria* at a relative abundance of 35% and 36% respectively. Similarly, the genus *Francisella* was also present at a relative abundance of 19% (*Theileria*-infected) and 10% (*Theileria*-uninfected) (Fig. 2C). Other genera found in *Theileria*-infected female *Hyalomma* ticks include the genus *Propionibacterium* (3.55%), *Bacillus* (12%), and *Acitenobacter* (6.3%). Interestingly, the genus *Ehrlichia* was found to be present in the uninfected female *Hyalomma* ticks at a relative abundance of (6.9%) (Fig. 2C). When compared to *A. marginale*-infected *Hyalomma* males, taxonomic abundance at the family level within the uninfected male *Hyalomma* ticks has a relative abundance of the family Propionibacteriaceae (29%) and Francisellaceae (12.7%). *A. marginale*-infected male *Hyalomma* ticks were predominantly dominated with bacteria within the family Anaplasmataceae (16.8%) and Staphylococcaceae (14.43%) (Fig. 1B). The bacteria family Candidatus Midichloriaceae and Francisellaceae were predominant in both *Theileria*-infected (35.3% and 18.9%) and uninfected (32.4% and 11.7%) *Hyalomma* ticks respectively (Fig. 2B).

**Figure 2:**
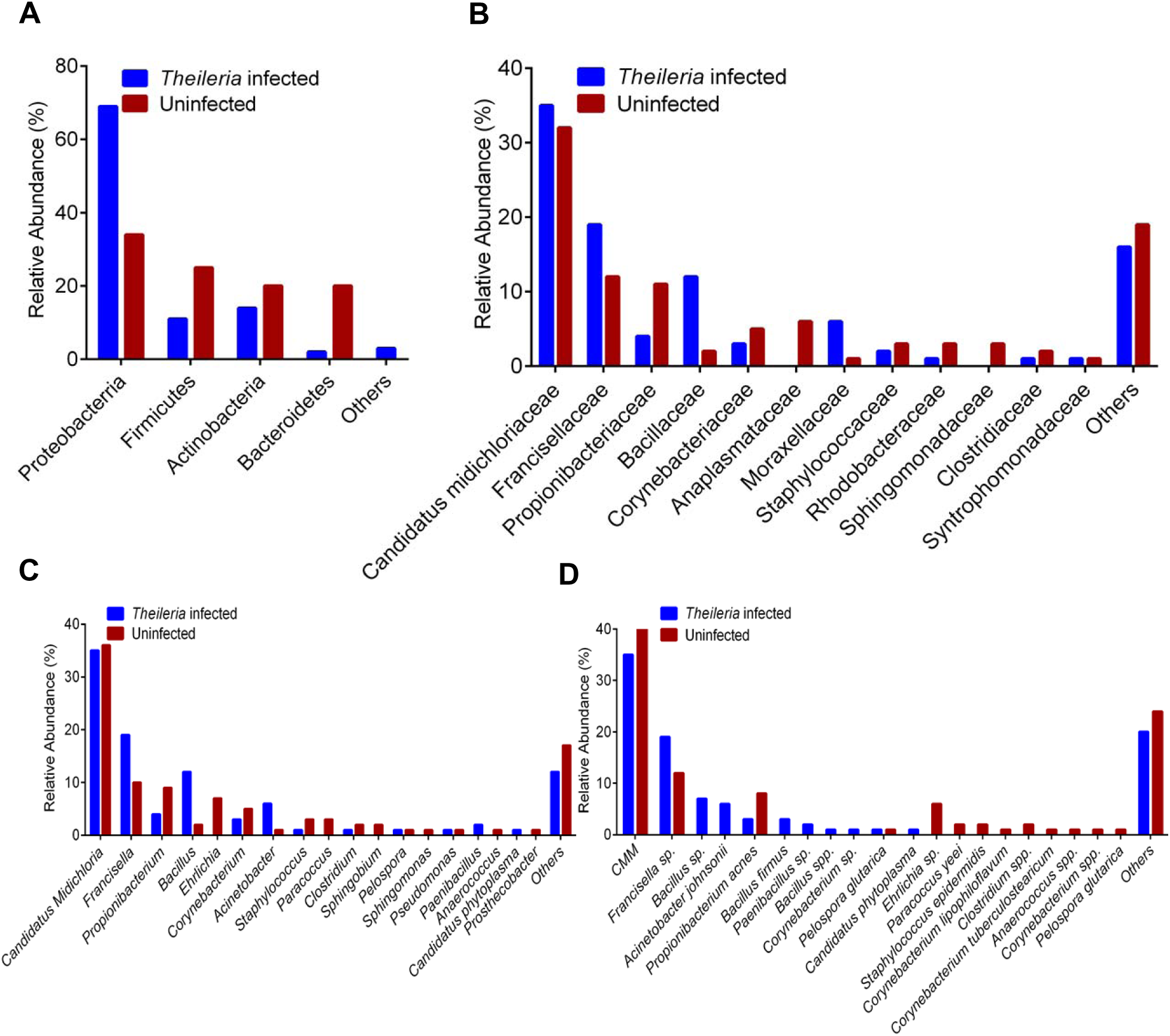
Relative abundance of taxa in *Theileria* species infected and uninfected female *H. anatolicum anatolicum* ticks at the A), phylum, B), family, C), genus and D), species level. The endosymbionts CMM and FLE were both detected in uninfected and *Theileria* species infected ticks. The species *Ehrlichia* was detected in the uninfected ticks.

Compared to the *Theileria* sp.-infected *Hyalomma* ticks, the family Anaplasmataceae (6.3%) and Propionibacteriaceae (11.2%) were more represented in the uninfected ticks (Fig. 2B). *A. marginale*-infected male *Hyalomma* ticks were also exclusively co-infected with *Ehrlichia* sp*., Anaplasma ovis*, *Staphylococcus (Staph.) sciuri*, *Acitenobacter (A.) indicus*, *Bacillus (Ba.) subtilis*, *Corynebacterium* sp., and *S. agnetis*. Interestingly, *Candidatus Midichloria mitochondrii* (CMM) and *Francisella* sp. were found to be present in both *A. marginale*-infected and uninfected male *Hyalomma* ticks albeit uninfected male *Hyalomma* having a higher abundance of *Francisella* sp. (Fig. 1D). Taxonomic distribution at the species level in the uninfected female *Hyalomma* ticks showed an exclusive presence of *Corynebacterium* sp*, Anaerococcus* sp*., Clostridium* sp*., Staph. epidermis,* and interestingly *Ehrlichia* sp., while *Bacillus* sp.*, Corynebacterium* sp*., Paenibacillus* sp., and *Ba. firmus* were exclusively found in the *Theileria* sp.-infected female *Hyalomma* ticks. While both infected and uninfected ticks both have CMM and *Francisella* sp. in their microbiome, uninfected ticks have a higher abundance of CMM, while infection with *Theileria* sp. was associated with increased the abundance of *Francisella* sp. (Fig. 2D).

Taxonomic distribution of Eukaryote revealed a higher abundance of the phylum Eukaryota in uninfected (67.57%) and *Theileria* sp-infected (51.50%) female *Hyalomma* ticks. The phylum Apicomplexa was also present in higher abundance in both groups (14.8% and 33.48%) respectively (Fig. 4A). At the species level, the microbial composition of uninfected female *Hyalomma* species was mostly dominated by *Gelidiella acerosa* (16.66%), *Trigonium formosum* (9.57%), *Phytophthora infestans* (8.62%) and *Hepatozoon (Hz.) americanum* (6.83%). Female *Hyalomma* ticks infected with *Theileria* sp. had a relative abundance of *Vaucheria canalicularis* (7.44%), *Plasmodium (P.) falciparum* (10.36%), and an increased abundance of *Hz. americanum* (14.9%) when compared to the uninfected ticks (Fig 4B).

#### Rhipicephalus microplus

The taxonomic abundance of microbial composition in uninfected *Rhipicephalus* ticks at the phylum level was represented by the phylum Proteobacteria (33.84%), Firmicutes (25.40%), Bacteroidetes (20.48%), and Actinobacteria (20.10%). The microbiome of *Theileria*-infected *R. microplus* was predominantly composed of bacteria from the phylum Firmicutes (96.8%) with the phyla Actinobacteria, Proteobacteria and Bacteroidetes also found to be present.

*A. marginale*-infected and uninfected *R. microplus* shared similar microbial composition. Flavobacteriaceae, Moraxellaceae, Staphylococcaceae, and Corynebacteriaceae were present at relatively similar distribution in both A*. marginale* infected and uninfected *R. microplus,* while the microbiome of *Theileria infected R. microplus* ticks was dominated by the bacteria from the family Bacillaceae (95.91%) (Fig. 3B).

**Figure 3:**
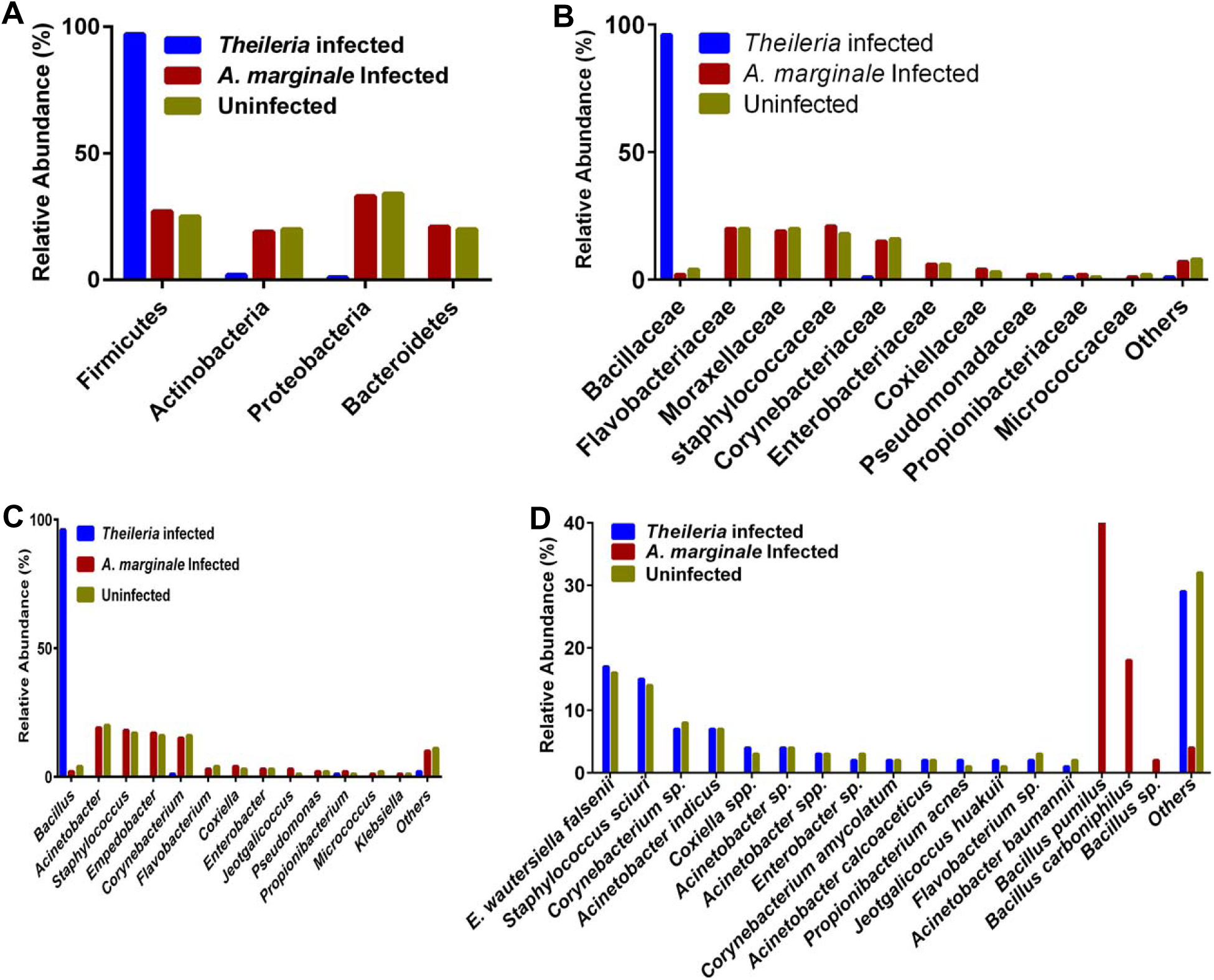
Relative abundances of taxa in *Theileria* infected, *A. marginale* infected and uninfected female *R. microplus* ticks at A), phylum, B), family, C), genus and D), species level. The phylum Firmicutes was the most abundant and three *Bacillus* species were detected at the species level in *Theileria* species infected *R. microplus* ticks.

Taxonomic abundance at the genus level showed *Bacillus* (95.72%) to be the most abundant within the *Theileria*-infected *R. microplus* ticks, while the composition of the clean and *A. marginale*-infected *R. microplus* ticks were closely related (Fig. 3C). *Bacillus* pumilus. (75.35%), *Ba. carboniphilus* (18.25%) and *Bacillus* sp. (1.94%) were the most abundant bacteria species present in *Theileria* sp.-infected *Rhipicephalus* ticks (Fig. 3D).

Uninfected and *A. marginale*-infected *R. microplus* ticks were both similarly represented at the species level albeit a slight increase in the abundance of *Coxiella* sp. from 2.96% in uninfected ticks to 4.25% in *A. marginale-*infected ticks (Fig. 3D). Distribution of Eukaryote at the phylum level was dominantly represented by the phylum Eukaryota (82.70%) in the uninfected *Rhipicephalus* ticks while the remaining phyla included Phaeophyceae (8.41%), Apicomplexa (4.822%), Euglenida (2.30%), and Xanthophyceae (1.65%). Uninfected and *A. marginale*-infected *R. microplus* had a relatively higher abundance of the phylum Eukaryota (45.07% and 77.53% respectively) (Fig. 4C). Species distribution of Eukaryote in *Rhipicephalus* ticks was also analyzed.

**Figure 4:**
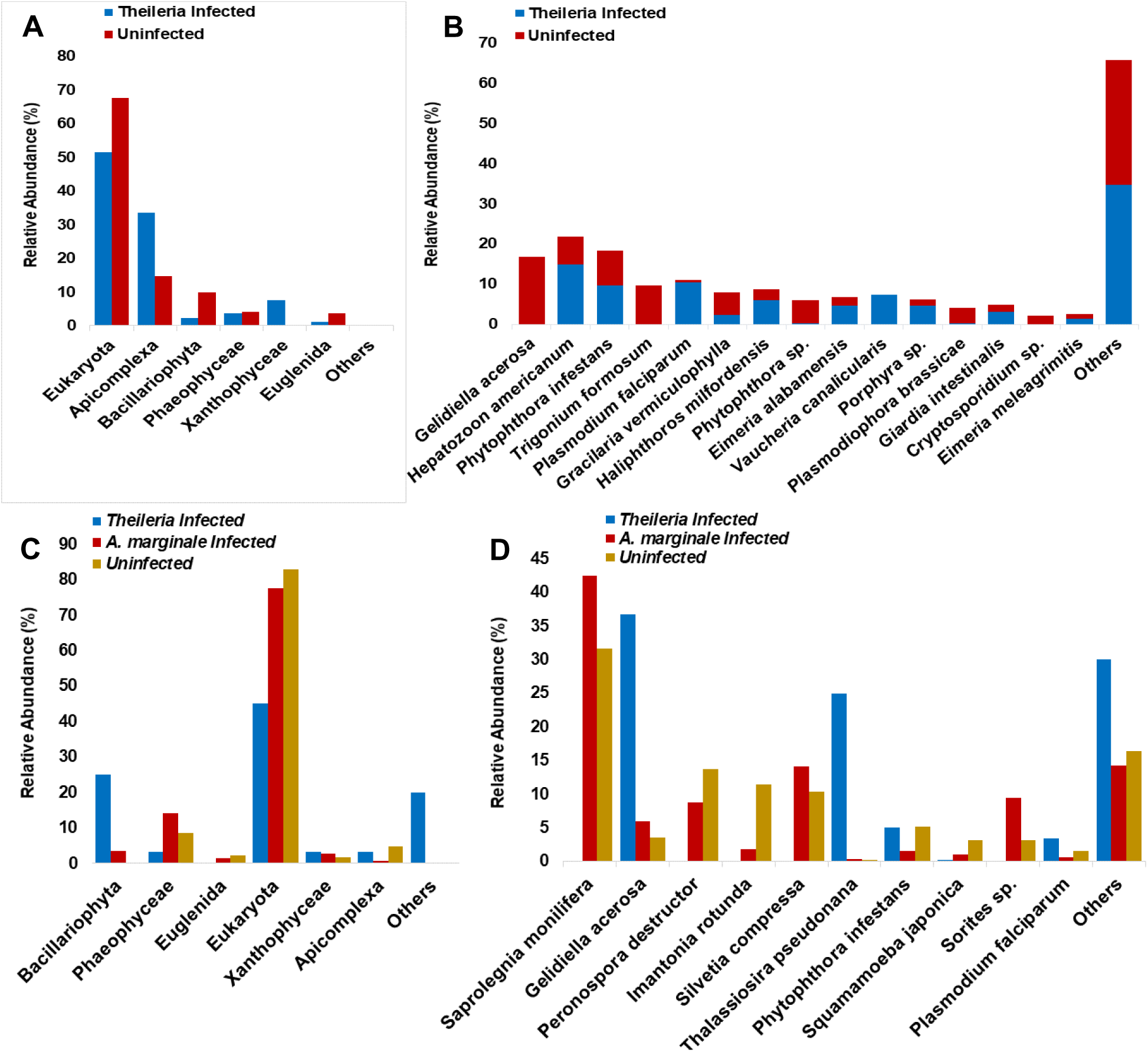
Relative abundance of eukaryote taxa. A and B represents phylum and species distribution in *Theileria* infected and uninfected female *H. anatolicum anatolicum* ticks respectively, while C and D represents phylum and species distribution in *Theileria* infected, *A. marginale* infected and uninfected female *R. microplus* ticks. *P. falciparum* and *Hz. americanum* were both detected from the 16S sequencing and both were relatively in higher abundance in *Theileria* infected ticks.

Both uninfected and *A. marginale*-infected *Rhipicephalus* ticks had relatively similar species distribution. Contrastingly, infection with *Theileria* sp. led to an increase in the abundances of *Gelidiella acerosa* (36.66%), *Thalassiosira pseudonana* (24.92%) and *P. falciparum* (3.33%) (Fig. 4D)

### Alpha Diversity

Why no significant differences was observed in the microbial richness between the two tick species (Faith_pd, p=0.858 and Observed OTUs, p=0.423), *H. anatolicum anatolicum* had a higher Faith_pd value and a higher number of OTU compared to *R. microplus* ticks (Fig. 6A and 6B). Pathogen infected ticks showed significant alpha-diversity based on Faith’s phylogenetic distance (p=0.0040) and number of observed OTUs (p=0.0040). *Theileria* sp.-infected *Hyalomma* ticks have a higher Faith_pd value and observed OTUs when compared to *Theileria* sp.-infected and *A. marginale*-infected *Rhipicephalus* ticks (Fig. 7A and 7B).

**Figure 5:**
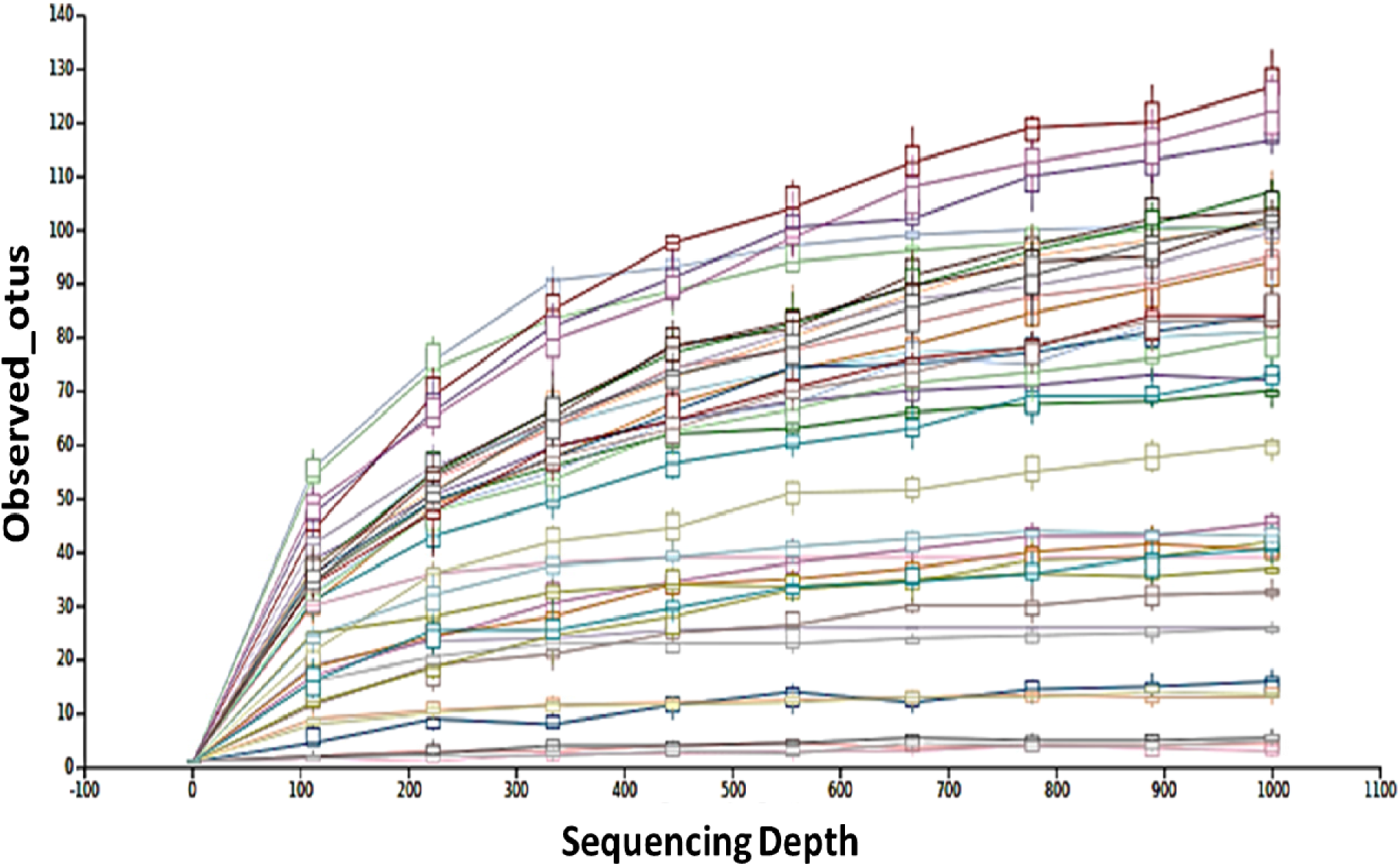
Rarefaction curves of individual tick samples rarefied to a sequence depth of 1000. This provides enough species diversity and allows for adequate sample sequence coverage for rarely existing or minimally represented operational taxonomic units. Ticks with less number of reads were removed from the analysis.

**Figure 6:**
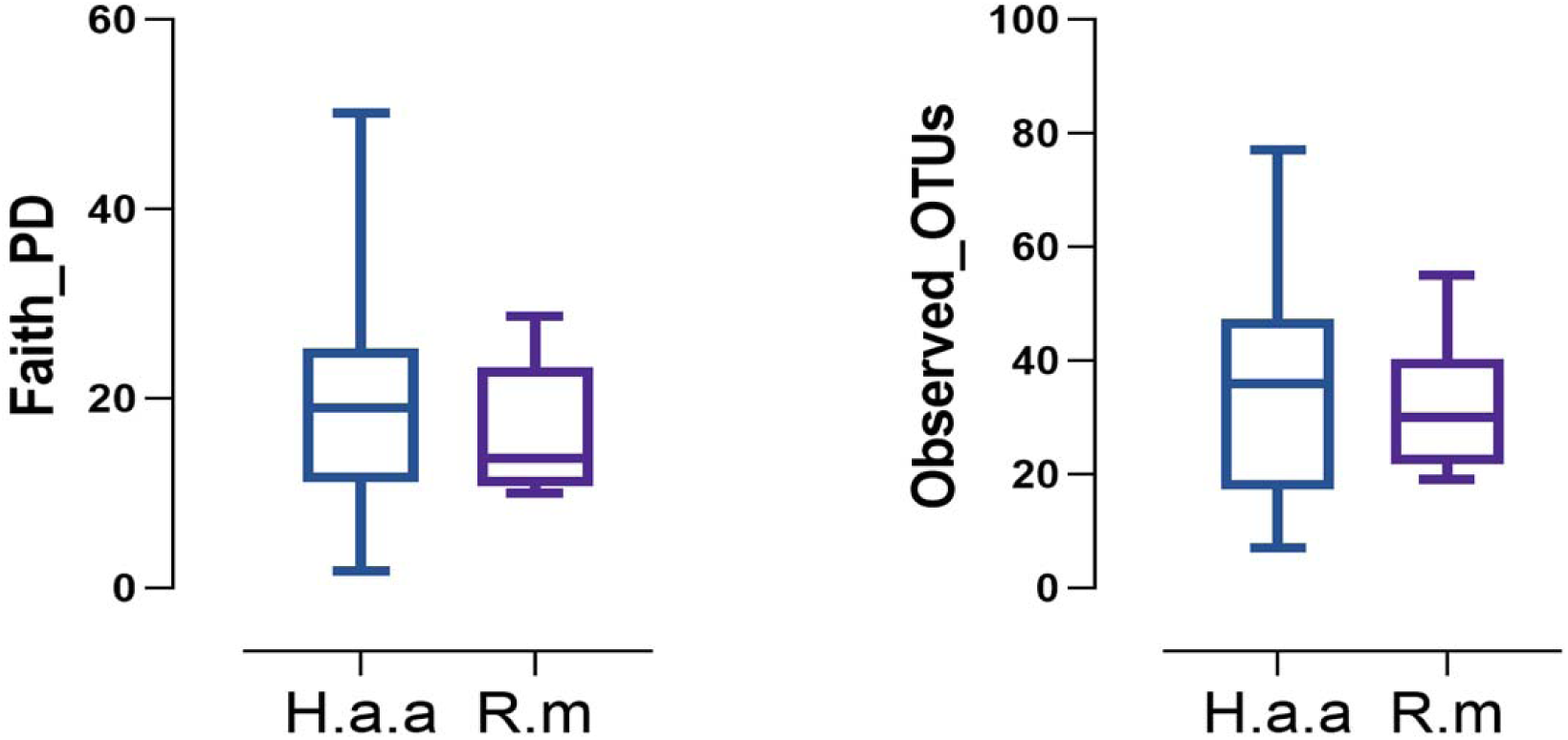
Alpha-diversity in uninfected *H. anatolicum anatolicum* (H.a.a) and *R. microplus* (R.m) ticks. Diversity metrics used for alpha diversity include A) Faith’s phylogenetic diversity, and B) observed OTUs. P > 0.05. *H. anatolicum anatolicum* ticks had a relatively more diverse microbiome with a higher OUT and Faith_pd value.

**Figure 7:**
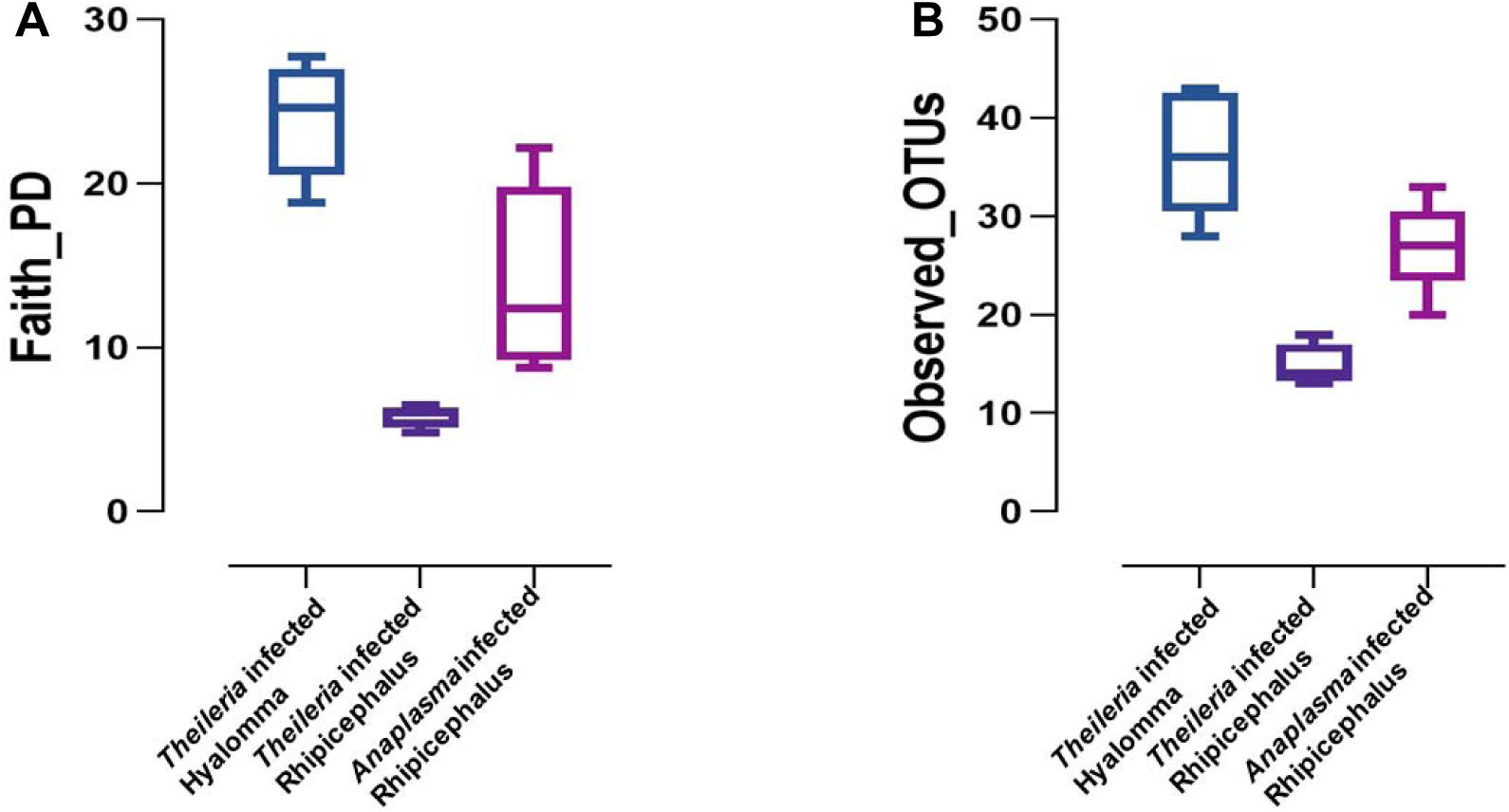
Alpha-diversity in infected *H. anatolicum anatolicum* (H.a.a) and *R. microplus* (R.m) ticks. Diversity metrics used for alpha diversity include A) Faith’s phylogenetic diversity, and B) observed OTUs. P < 0.05. *Theileria* infected *R. microlplus* ticks was the least diverse in contrast to *Theileria* infected *H. anatolicum anatolicum* which had the most diverse microbiome.

### Genetic relationship of selected *Coxiella*, *Anaplasma*, and *Ehrlichia* sequences

The two *Coxiella* sequences from this study uniquely grouped with a *Coxiella burnetii* sequence (GenBank: NR 104916.1)

### Beta Diversity

Principal coordinate analysis (PCoA) of the distance matrixes showed clustered separation between *Hyalomma a. anatolicum* and *R. microplus* ticks (Fig 8A and 8B). Distinct clustering was also observed on the PCoA plot of infected ticks using the unweighted and weighted distance matrices (Fig 9A and 9B). Additional information can be seen on the supplementary data (S1; Table 2 and 3)

**Figure 8:**
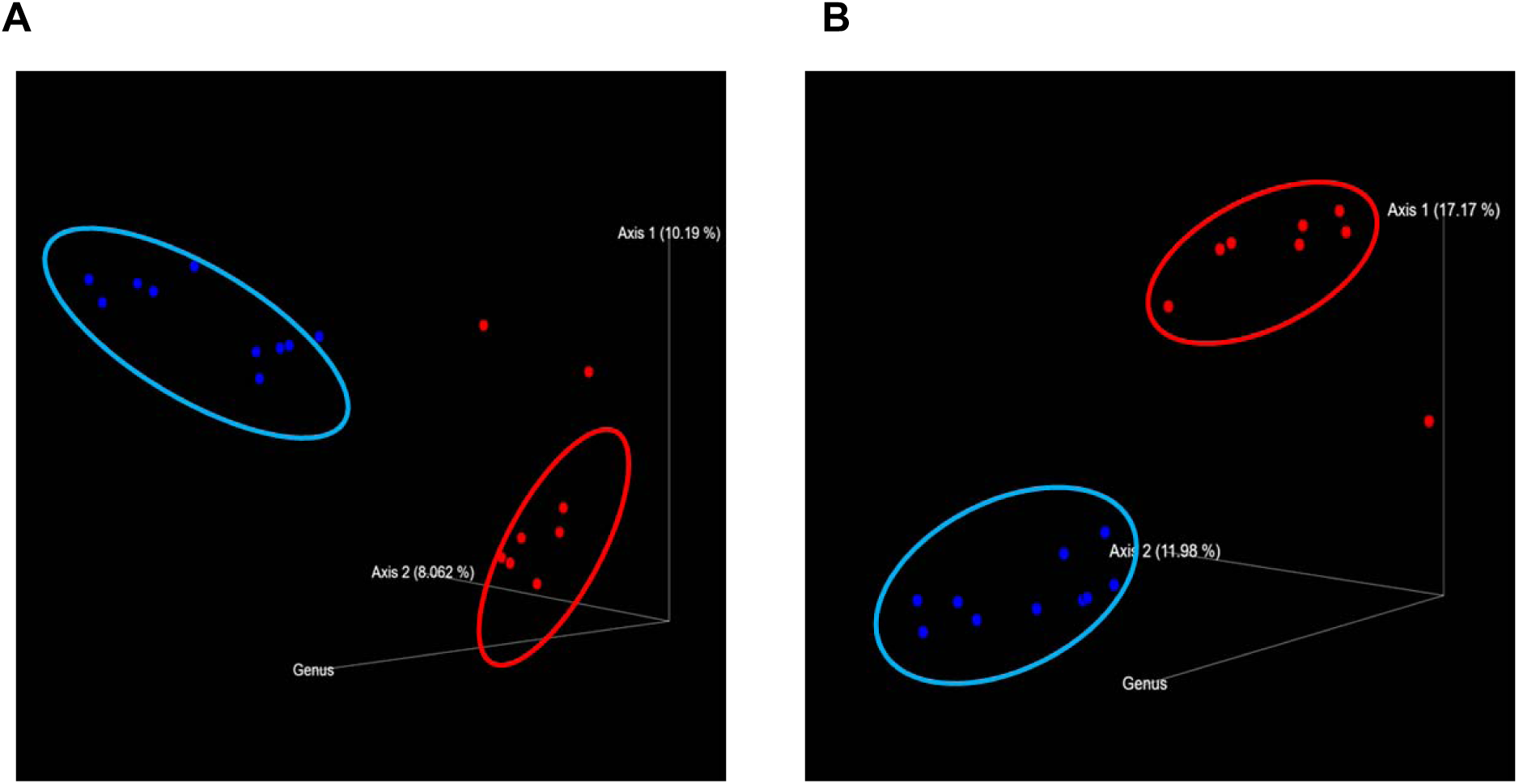
Principal coordinate analysis (PCoA) plots of uninfected ticks using A), unweighted and B), weighted UniFrac distance matrices. Red points and circles represents *H. anatolicum anatolicum* and *R. microplus* is represented in blue. Colored points represents individual biological replicates. Each tick species shows distinct clustering from one another with few outliers seen in the *Hyalomma* ticks.

**Figure 9:**
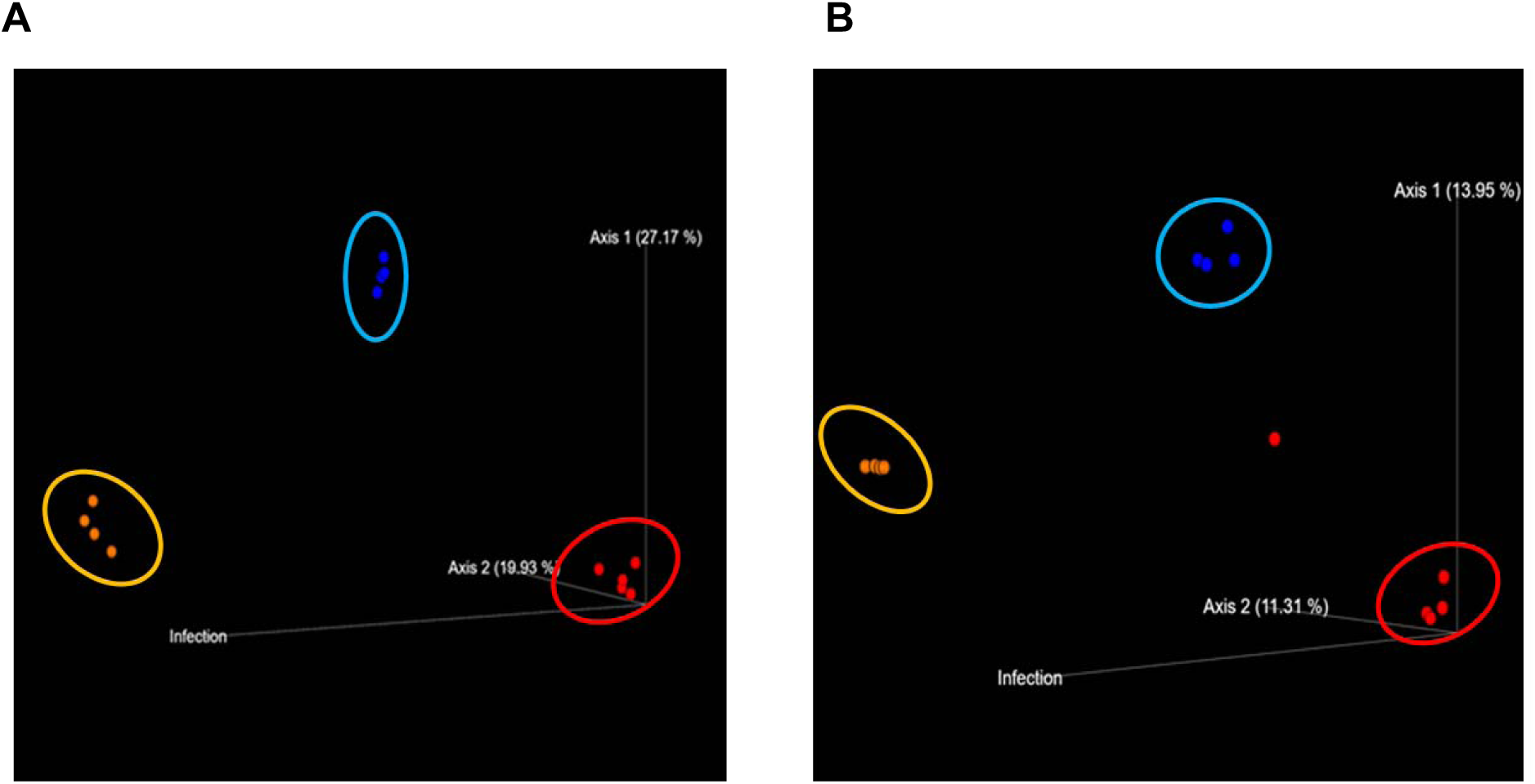
Principal coordinate analysis (PCoA) plots of *Theileria* infected *H. anatolicum anatolicum* (red), *Theileria* infected *R. microplus* (blue) and *A. marginale* infected *R. microplus* (yellow) ticks using A), unweighted and B), weighted UniFrac distance matrices. Clustered separation of the ticks a result of the pathogen presence shows that all three groups shares little to no similar microbial composition.

## Discussion

To the best of our knowledge, this study is the first to evaluate the microbiome composition of pathogen-infected and uninfected *H. anatolicum anatolicum* and *R. microplus* ticks infesting livestock from Pakistan. While ticks from the genus *Hyalomma* and *Rhipicephalus* have previously been reported to harbor both protozoan and bacterial organisms that can potentially be transmitted to their livestock hosts during feeding (9–10, 17, 53), this study expands on existing knowledge of the microbial communities residing within the tick vector and the possible interactions with their associated pathogens. Likewise, previous microbiome studies of ticks from Pakistan, have employed the 454 pyrosequencing techniques (10, 12), for this study, we utilized a high throughput Illumina sequencing approach for microbial analysis. Similar studies of microbiome of hard ticks in general have also employed high throughput Illumina sequencing approach, (13–16), few of those only investigated the microbial patterns in infected and uninfected ticks (16).

In this study, the prevalence of *Theileria* species as seen in previous studies, was higher in *H. anatolicum anatolicum* ticks, while *A. marginale* had similar prevalence in both tick species (7, 9, 17, 18) (Table 1). With the exception of the *R. microplus* ticks, microbiome composition at the phylum taxonomic level was predominantly dominated by the phylum Proteobacteria in all tick groups tested in this study (Fig 1A and 2A). Previous studies of tick microbiome composition have reported a dominant abundance of the member of Phylum Proteobacteria (12–13, 21–22), one of the most diverse bacterial group which includes free-living commensals and pathogenic species that infect humans and animals alike.

Male *H. anatolicum anatolicum* ticks infected with *A. marginale* were also found to be co-infected with bacteria in the genera *Anaplasma* and *Ehrlichia*. This finding raised important questions concerning the competence of the male ticks in pathogen transmission as these genera are of pathogenic significance. While male ticks have not been associated with natural pathogen transmission, an elegant study by Zivkovic *et al* demonstrates vector competence of *Dermacentor reticulatus* for *A. marginale* (23).

The ability of two or more pathogen to co-infect male ticks with subsequent transmission to the host still needs to be investigated, the genera *Anaplasma* and *Ehrlichia*, both of which are *Anaplasmataceae* and obligate intracellular pathogens could both occur synergistically thus favoring co-existence within the same tick vector. Another source of the coinfection could be from the blood of the host animal the ticks were feeding on as at the time of collection.

We also revealed the *Francisella* genus in uninfected male *H. anatolicum anatolicum* ticks. This is possibly an endosymbiont that has also been previously reported in other tick species and has been shown to have parts of its genome that encodes for biosynthesis of B vitamins (24), hence helping the tick in compensating for the nutrient poor blood meal. Since male ticks rarely spend lesser time blood-feeding when compared to their female counterparts, in the presence of *Francisella,* it could be hypothesized that male ticks also utilize the B vitamins for reproductive development such as reproductive hormone production and nutrition of the sperm cells. Since male ticks have also been shown to be early arrivers for a blood meal in order to release pheromones that will subsequently attract females, this process could place a lot of physiological stressors (25, 26) on the male ticks hence the maintenance of the *Francisella* genus for its B vitamins synthesis.

The endosymbionts CMM and (FLE) were both detected in uninfected and *Theileria* sp.-infected *H. anatolicum anatolicum* ticks (Fig 2C and 2D). These are obligate, vertically maintained endosymbionts in the phylum Proteobacteria. While they have been hypothesized to nutritionally support the tick host, their role in pathogen acquisition, susceptibility and transmission within their tick host is still yet to be explained. *Rickettsia parkeri*, a spotted fever group rickettsia transmitted by *Amblyomma maculatum* ticks was shown to selectively reduce the abundance of FLE while favoring an increase in CMM (27–28). A similar interaction was observed in our study as *Theileria* sp.-infected female *H. anatolicum anatolicum* ticks have a higher abundance of FLE as compared to uninfected ticks that has more CMM in their microbiome. While the FLE has been reportedly found in the genus *Hyalomma* (28–30), to the best of our knowledge, this is the first study to report CMM and FLE in *H. anatolicum anatolicum* ticks from Pakistan.

We also observed interactions at the species level with or without pathogen infection. Of note were male ticks that were PCR positive with *A. marginale* were also exclusively infected with *A. ovis*, *Ehrlichia* sp., *Acinetobacter indicus*, and *Bacillus subtilis*. Similarly, *Theileria* sp. infection in female *R. microplus* was found to have an abundance of *Bacillus* sp., *Ba. carboniphilus,* and *Ba. firmus.* The genus *Bacillus* belongs to the phylum Firmicutes and the species are known for their ability to form endospores in unfavorable environmental conditions. Their relative abundance in *A. marginale*-infected male *H. anatolicum anatolicum* and *Theileria*-infected *R. microplus* ticks could be due to a pathogen-associated dysbiosis of the microbial community within the ticks. The success of a pathogen to successfully colonize within the tick host depends partly on the pathogen’s ability to disrupt the existing microbial community normally present within the tick. Colonization by *Theileria* sp. and *A. marginale* could cause a pathogen-induced dysbiosis which could be by their metabolic activities in which they produce toxic metabolites or compete for available nutrients. Since Bacilli are spore forming bacteria, this will make them at an advantage to survive in extremely unfavorable conditions, which could be a reason for their increased abundance in the infected ticks.

*Coxiella* sp. was also detected in uninfected and *A. marginale*-infected female *R. microplus* ticks albeit at a relatively higher abundance in the infected ticks. A previous research has reported *Coxiella* sp. in *R. microplus* ticks (1). The *Coxiella* sp. reported in this study is likely a Coxiella-Like Endosymbiont (CLE) which has been previously reported in other tick species (5, 31–32) and its relationship in tick has also been associated with the survival of their tick host, where the CLE genome has been shown to encode biosynthesis pathways of major B vitamins (B7, B9 and B12), nutrients which are deficient in the blood meal (33–34). The increased *Coxiella* sp. abundance in *A. marginale*-infected ticks could be attributed to a possible dependency of *A. marginale* on B vitamins for its lifecycle within the tick host. As both *A. marginale* and *Coxiella* sp. are obligate and intracellular in nature, it is safe to suggest that *A. marginale*, in trying to evade the tick’s innate immune response will hide behind the self-recognition of CLE by the tick’s immune cells, hence becoming an effective pathogen.

The most abundant bacteria species in uninfected and *A. marginale*-infected *R. microplus* ticks were *Empedobacter wautersiella, Staph. sciuri, Corynebacterium sp., Acinetobacter indicus, Coxiella* sp.*, Acinetobacter sp.,* and *Enterobacter* sp. The maintenance of these bacterial species in both uninfected and *A. marginale*-infected *R. microplus* ticks could be due to the fact that some of these bacteria are important in the tick biology. *Acinetobacter* sp. and *Flavobacterium* sp., though with unclear roles to the arthropod hosts, have been shown to increase the growth rate and the survival of stable fly larvae (35), while *Acitenobacter* has also been reported in *Ixodes ricinus* (36) and some blood feeding arthropods (37–38). Surprisingly, we observed a complete shift in the bacterial species composition in the microbial composition of *Theileria* infected *R. microplus.* Approximately 96% of the bacterial species belongs to the genus *Bacillus* with *Ba. pumilus*, *Ba. carboniphilus* and *Bacillus* sp. been the most predominant (Fig. 7A). While studies have shown how *Theileria* pathogens interfere with the mammalian host’s immune response by expressing proteins necessary for its transformation (51), its impact on the tick microbiota has yet to be shown. Inhibition of important microbial metabolic pathways by *Theileria*-associated proteins (52) could have led to a pathogen-associated dysbiosis.

The endosymbionts CMM and FLE were both detected in uninfected and *Theileria*-infected female *H. anatolicum anatolicum* at relatively unchanged abundances (Fig. 2A). Although direct competition for nutrient and niche between endosymbionts and pathogen has been reported to take place in ticks (6), our finding did not support this as the abundances of these endosymbionts were not altered in the presence or absence of *Theileria* infection. The reason for this could arise from the differences in the routes by which the endosymbionts and *Theileria* are maintained within the tick vector. As obligate endosymbionts, both CMM and FLE have been shown to be transovarially maintained from the female ticks to the eggs (27), while *Theileria* is transtadially maintained. This difference in their trafficking within the tick vector could lead to a reduced or no chance of them interacting together within the tick.

While alpha diversity analysis of uninfected *H. anatolicum anatolicum* and *R. microplus* ticks suggested no significant differences in the bacterial richness (Fig. 6A and B), cluster analysis revealed that the two ticks have unrelated microbial composition based on the number of OTUs and the phylogenetic distances of individual bacterial species (Fig 8A and B). This finding was further supported by the differences in the bacterial composition at the genus level (Fig 2C and 3C). This difference in the microbial composition could be as a result of their feeding habit, *Hyalomma* ticks are known to be two/three-host ticks while *Rhipicephalus* ticks are generally a one-host tick. Unsurprisingly, significant differences were seen in the alpha diversity analyses of pathogen infected ticks. *Theileria*-infected *R. microplus* ticks were the least diverse with a lesser amount of OTUs as well as a relatively low Faith_pd value (7A and 7B). This further supported our previous pathogen-associated dysbiosis that was seen in *Theileria*-infected *R. microplus* ticks (Fig 3D).

Other bacteria species found to be abundant in *Theileria*-infected *H. anatolicum anatolicum* were *Bacillus sp., Acinetobacter johnsonii, Propionibacterium (Pr.) acnes and Bacillus firmus*, while uninfected ticks had an abundance of *Pr. acnes* and *Ehrlichia sp*. Surprisingly, apicomplexan pathogen *Hz. americanum* and *P. falciparum* were detected in both *R. microplus* and *H. anatolicum anatolicum*. *Hz. americanum*, the cause of American canine hepatozoonosis (ACH) an emerging disease of dogs has been previously reported in Taiwan and Thailand where transmission has been shown to be by *R. sanguineus* (40–41). In the United States, it is vectored by the *Amblyomma maculatum* nymphs or adults (39).

Pakistan is a malaria endemic country and we observed an increase in the relative abundance of *P. falciparum* in *Theileria*-infected *H. anatolicum anatolicum* when compared to the uninfected ticks (Fig. 5A). Similar observation was made in *Theileria*-infected *R. microplus* ticks (Fig 8A). The detection of *P. falciparum* the protozoan parasite, a causative agent of the human malaria, in ticks was an unaccepted finding in this study. The detection of *P. falciparum* in ticks from this study could have risen from the ticks accidentally feeding on an infected human host. While this is a possibility in a multi-host ticks as seen in *Hyalomma* species, this is highly an unlikely occurrence in *R. microplus* which is a one-host tick. The presence of *P. falciparum* in both *R. microplus* and *H. anatolicum anatolicum* warrants further investigation. To our knowledge, ticks are neither competent vector nor reservoir of *P. falciparum* and further studies are needed to understand the presence of *P. falciparum* in tick species.

## Conclusions

*Empedobacter wautersiella, Staph. sciuri, Corynebacterium sp., Acinetobacter indicus, Coxiella* sp.*, Acinetobacter* sp., and *Enterobacter* sp. were detected in abundance in uninfected and *A. marginale* infected *R. microplus* while about 95% of the bacteria species in the *Theileria*-infected *R. microplus* ticks belong to the genus *Bacillus*. CMM and FLE were both detected in abundance (>50%) in the microbiome of uninfected and *Theileria*-infected *H. anatolicum anatolicum* ticks. Our study revealed that the infection of tick with a protozoan pathogen leads to dysbiosis of the bacterial community and this was seen in the *Theileria*-infected *R. microplus* ticks where *Bacillus firmus*, *Ba. carboniphilus and Bacillus sp* represented about 95% of the total microbiome. This observation was further confirmed by a very low number of operational taxonomic unit (OTUs) and Faith’s phylogenetic distance value, both of which are measures of species richness. In contrast to the bacterial dysbiosis seen in *Theileria*-infected *R. microplus ticks, Theileria*-infected female *H. anatolicum anatolicum* ticks were co-infected with the apicomplexan parasites *P. falciparum* and *Hz. americanum*. We also showed that infection of tick with a bacteria pathogen do not lead to a change in the microbiome of the tick vector, though it could predispose the tick to been co-infected with other pathogenic bacterial species as shown by the co-infection of *A. marginale* infected male *H. anatolicum* ticks with *Ehrlichia sp* and *A. ovis*. In all, *Theileria* infection in the ticks reduces bacterial diversity and was also found co-infecting the tick alongside other apicomplexan pathogen, while *A. marginale* infection was correlated *A. ovis* and *Ehrlichia sp* co-infection.

This study establishes the extent of the diversity of microbial community within two important tick species from Pakistan and revealed the presence of *Theileria*, *A. marginale* and additional pathogenic bacteria that could be of public health significance. We hypothesized that infection with either a protozoan or bacteria pathogen will alter the microbial composition within these tick specie. Limitation faced during this study was the difficulty in getting tissue samples of ticks as they were field collected. Future tick developmental and tissue-specific studies warrant new insights in specific interactions between tick-borne pathogens and their associated microbiome.

## Methods

### TICK COLLECTION

*H. Anatolicum anatolicum* and *R, microplus* ticks were carefully removed from cattle, sheep and goats from Sialkot [32°29′33.7″N, 74°31′52.8″E], Gujrat [32°34′22″N, 74°04′44″ E], Gujranwala [32°9′24″N, 74°11′24″E], and Sheikhupura [31°42 47″ N, 73°58′41″ E] districts located in the province of Punjab, Pakistan. Briefly, fully engorged ticks were carefully removed from the body of the animals with the mouth part intact using tweezers. All ticks were kept in separate vials containing 70% ethanol and details of the location, sex, and host were recorded. For this study, a total of 320 ticks were selected and shipped from Pakistan to the University of Southern Mississippi for further analysis using the U.S. Department of Agriculture’s Animal and Plant Health Inspection Service (permit # 11122050).

### Tick species identification

Tick identification was performed by an expert taxonomist (Dmitry A. Apanaskevich) at the United States National Tick Collection (USNTC) according to the criteria used in previously published reports (42–44). All stages were examined on an Olympus SZX16 stereoscopic microscope.

### Genomic DNA Extraction

Prior to DNA extraction, individual ticks were surface sterilized in a series of steps. Briefly, a 10% solution of sodium hypochlorite was used to clean the individual surfaces of ticks followed by rinsing in 70% ethanol. A final cleaning was done using sterile water. Genomic DNA was extracted from each individual tick homogenate using a DNeasy blood and tissue kit (Qiagen, Valencia, CA, USA) following the manufacturer’s protocol. The concentrations of the extracted genomic DNA samples were quantified using a Nanodrop ND-100 instrument and DNA stored in −20°C till further needed.

### Detection of *Theileria* species

To detect *Theileria* species from the collected ticks, forward and reverse primers specific for the *Theileria* genus 18S rRNA gene were used in a 25 μL reaction volume (10). The reaction volume consisted of 50-75 ng of genomic DNA, 1 μL each of both primers, 12.5 μL of PCR 2X Master Mix (Biolab Inc.) and the remainder nuclease free water. The reaction mixture was subjected to thermal cycling at 94°C for 3 min followed by 39 cycles of 94°C for 20 s, 48°C for 60 s, and 68°C for 30 s, and a final extension step at 68°C for 2 min. 8 μL of the amplified product were electrophoresed in an ethidium bromide stained 2% gel. The amplicons obtained were isolated and purified using a QIAquick PCR purification kit (Qiagen), and the purified products were sequenced by Eurofins. The partial sequences obtained were subjected to the NCBI BLAST program for species identification of the *Piroplasma* sequences.

### Detection of *Anaplasma marginale*

The detection of *Anaplasma marginale* was carried out as previously described (45). *Anaplasma marginale* 16S rRNA forward Amar16S-F: GGC GGT GAT CTG TAG CTG GTC TGA and reverse primers Amar16S-R: GCC CAA TAA TTC CGA ACA ACG CTT were used in a 25 μL reaction volume containing 1μL each of both primers, 50-70 ng of DNA, 12.5 μL of PCR 2X Master Mix (Biolab Inc.) and nuclease free water to make the final volume. The reaction mixture was subjected to thermal cycling at 94°C for 5 min followed by 35 cycles of 94°C for 45 s, 55°C for 45 s, and 72°C for 45 s, and a final extension step at 72°C for 10 min. 8 μL of the amplified product were electrophoresed in an ethidium bromide stained 2% gel The amplicons obtained were isolated and purified using a QIAquick pcr purification kit (Qiagen), and the purified products were sequenced by Eurofins. The partial sequences obtained were subjected to the NCBI BLAST program for species identification of the *Anaplasma* sequences.

### 16S rRNA Library Preparation and Sequencing

A total of 40 ticks representing 20 *H. anatolicum anatolicum* and 20 *R. microplus* ticks were divided into 5 biological replicates depending on the presence or absence of *Theileria* and *Anaplasma marginale* infection. The *Hyalomma* ticks comprises of 5 males and 15 female ticks, while the *Rhipicephalus* ticks are all females. PCR sequencing of the V1-V3 variable region of the bacterial 16S rRNA gene were amplified using barcoded primers 27F/519R as outlined by the 16S Illumina’s MiSeq protocol (www.mrdnalab.com, Shallowater, TX, USA). Briefly, PCR was performed using the HotStarTaq Plus Master Mix Kit (Qiagen, USA) under the following conditions: 94°C for 3 min, followed by 30-35 cycles of 94°C for 30 s, 53°C for 40 s and 72°C for 1 min, after which a final elongation step at 72°C for 5 min was performed. After amplification, PCR products were electrophoresed in 2% agarose gel to determine the success of amplification and the relative intensity of bands. Multiple samples are pooled together in equal proportions based on their molecular weight and DNA concentrations. Pooled samples are purified using calibrated Ampure XP beads. Then the pooled and purified PCR product is used to prepare Illumina DNA library. Sequencing was performed at MR DNA (www.mrdnalab.com, Shallowater, TX, USA) on a MiSeq following the manufacturer’s guidelines.

### Phylogenetic Analyses

The *Anaplasma*, *Ehrlichia* and *Coxiella* partial 16S rRNA sequences were obtained and genetic relationships was compared with similar sequences from NCBI. Briefly, downloaded reference sequences were downloaded, aligned using the ClustalW and phylogenetic tree was constructed (S1, Fig. 1)

The evolutionary history was inferred using the Neighbor-Joining method (56). The bootstrap consensus tree inferred from 1000 replicates (57) is taken to represent the evolutionary history of the taxa analyzed (57). Branches corresponding to partitions reproduced in less than 50% bootstrap replicates are collapsed. The percentage of replicate trees in which the associated taxa clustered together in the bootstrap test (1000 replicates) are shown next to the branches (57). The evolutionary distances were computed using the p-distance method (58) and are in the units of the number of base differences per site. This analysis involved 25 nucleotide sequences. Codon positions included were 1st+2nd+3rd+Noncoding. All ambiguous positions were removed for each sequence pair (pairwise deletion option). There were a total of 1512 positions in the final dataset. Evolutionary analyses were conducted in MEGA X (59).

### Data Analyses

Sequence analysis was done using the Quantitative Insights into Microbial Ecology (QIIME 2) (46), unless stated otherwise. Briefly, processing of raw FASTQ files were demultiplexed. The Atacama soil microbiome pipeline was incorporated for quality control of demultiplexed paired-end reads using the DADA2 plugin as previously described (47). Sequence alignment and subsequent construction of phylogenetic tree from representative sequences was done using the MAFFT v7 and FasTree v2.1 plugin (48). Operational taxonomic assignment was done using the qiime2 feature-classifier plugin v7.0 (49) which was previously trained against the SILVA 132 database preclustered at 99% (50).

Diversity analysis was done by rarifying individual sequences to a depth of 1000 to get adequate coverage of all samples analyzed and samples with insufficient reads subsequently screened out (Fig. 5). Diversity analysis was only estimated for female tick samples as the number of male ticks were small to give a confident result. Faith phylogenetic distance (Faith_pd) and the number of observed OTUs were used in assessing alpha diversity, while beta diversity was estimated using PERMANOVA analysis of unweighted and weighted UniFrac distance matrix. Raw data from this analysis were submitted deposited and assigned the GenBank BioProject number PRJNA600935.

## Acknowledgements

This research was principally supported by a Pakistan-US Science and Technology Cooperation Program award (US Department of State) and the Mississippi INBRE, funded by an institutional Award (IDeA) from the National Institute of General Medical Sciences of the National Institutes of Health under award # P20GM103476. The funders played no role in the study design, data collection and analysis, decision to publish, or preparation of the manuscript.

## Competing interests

The authors have declared that no competing interests exist.

## Authors Contributions

Conceptualization: Shahid Karim

Data curation: Abdulsalam Adegoke, Deepak Kumar, Shahid Karim

Formal analysis: Abdulsalam Adegoke, Deepak Kumar, Shahid Karim

Funding acquisition: Shahid Karim, Muhammad Imran Rashid, Aneela Zameer Durrani, Muhammed Sohail Sajid

Investigation: Abdulsalam Adegoke, Deepak Kumar, Muhammad Imran Rashid, Aneela Zameer Durrani, Muhammed Sohail Sajid, Shahid Karim

Methodology: Abdulsalam Adegoke, Shahid Karim

Project administration: Shahid Karim, Muhammad Imran Rashid, Aneela Zameer Durrani, Muhammed Sohail Sajid

Resources: Shahid Karim, Muhammad Imran Rashid, Aneela Zameer Durrani, Muhammed Sohail Sajid

Supervision; Shahid Karim

Validation: Abdulsalam Adegoke, Deepak Kumar, Shahid Karim

Visualization: Abdulsalam Adegoke, Deepak Kumar, Shahid Karim

Writing-original draft: Abdulsalam Adegoke, Shahid Karim

Writing-review & editing: Shahid Karim

## LIST OF SUPPLEMENTARY TABLES

**S1; Table 5:** Species identification of *Theileria* species following sequencing

| ***Blast result*** | ***% Identity*** | ***Accession*** |
| --- | --- | --- |
| *Anaplasma marginale* | 90.80% | KU586030.1 |
| *Anaplasma marginale* | 96.43% | JF949767.1 |
| *Anaplasma marginale* | 90.00% | KU585987.1 |
| *Anaplasma marginale* | 93.21% | JF949767.1 |
| *Anaplasma marginale* | 91.77% | MK016525.1 |
| *Anaplasma marginale* | 91.77% | CP023731.1 |
| *Anaplasma marginale* | 91.77% | CP023730.1 |
| *Anaplasma marginale* | 91.77% | FJ155998.1 |
| *Anaplasma marginale* | 92.41% | MK016525.1 |
| *Theileria orientalis* | 99.93% | AP011949.1 |
| *Theileria lestoquardi* | 96.50% | MG208059.1 |
| *Theileria annulata* | 95.98% | MK415835.1 |
| *Theileria annulata* | 95.74% | MK918607.1 |
| *Theileria annulata* | 95.74% | MK415058.1 |
| *Theileria annulata* | 95.74% | MK838106.1 |
| *Theileria annulata* | 95.74% | MK183002.1 |
| *Theileria annulata* | 95.74% | MK183000.1 |
| *Theileria annulata* | 95.74% | MK182999.1 |
| *Theileria annulata* | 95.74% | MK182996.1 |
| *Theileria annulata* | 100.00% | KT004406.1 |
| *Theileria annulata* | 95.74% | MN625889.1 |
| *Theileria annulata* | 95.74% | MN625888.1 |
| *Theileria equi* | 100.00% | KY464022.1 |
| *Theileria equi* | 98.00% | KY464019.1 |
| *Theileria equi* | 98.17% | KY464020.1 |
| *Theileria equi* | 97.67% | MG569904.1 |
| *Theileria equi* | 97.61% | JX177673.1 |
| *Theileria equi* | 97.55% | KX227640.1 |
| *Theileria equi* | 97.55% | MK615933.1 |
| *Theileria equi* | 97.55% | JX177671.1 |
| *Theileria equi* | 100.00% | MK346272.1 |

**S1; Table 2:** Beta-diversity metrics from uninfected *H. anatolicum anatolicum* and *R. microplus* ticks

|  | Group 1 | Group 2 | Sample size | Permutations | pseudo-F | p-value | q-value |
| --- | --- | --- | --- | --- | --- | --- | --- |
| Bray_Curtis | *H. anatolicum anatolicum* | *R. microplus* | 18 | 999 | 3.197098313 | 0.001 | 0.001 |
| Jaccard | *H. anatolicum anatolicum* | *R. microplus* | 18 | 999 | 1.629955018 | 0.001 | 0.001 |
| unweghted_UniFrac | *H. anatolicum anatolicum* | *R. microplus* | 18 | 999 | 3.028314007 | 0.001 | 0.001 |
| weighted_UniFrac | *H. anatolicum anatolicum* | *R. microplus* | 18 | 999 | 11.11746648 | 0.001 | 0.001 |

**S1; Table 4:** Beta-diversity metrics from *Theileria* species and *A. marginale* infected *H. anatolicum* and *R. microplus*.

|  | Group 1 | Group 2 | Sample size | Permutations | pseudo-F | p-value | q-value |
| --- | --- | --- | --- | --- | --- | --- | --- |
| Bray_Curtis | *A. marginale* infected *R. microplus* | *Theileria* infected *H. anatolicum* | 10 | 999 | 2.051402834 | 0.027 | 0.027 |
|  | *A. marginale* infected *R. microplus* | *Theileria* infected *R. microplus* | 9 | 999 | 1.804879723 | 0.018 | 0.027 |
|  | *Theileria* infected *H. anatolicum* | *Theileria* infected *R. microplus* | 9 | 999 | 2.688977673 | 0.02 | 0.027 |
| Jaccard | *A. marginale* infected *R. microplus* | *Theileria* infected *H. anatolicum* | 10 | 999 | 1.508145443 | 0.005 | 0.0075 |
|  | *A. marginale* infected *R. microplus* | *Theileria* infected *R. microplus* | 9 | 999 | 1.659339366 | 0.005 | 0.0075 |
|  | *Theileria* infected *H. anatolicum* | *Theileria* infected *R. microplus* | 9 | 999 | 1.992545142 | 0.01 | 0.01 |
| unweighted_UniFrac | *A. marginale* infected *R. microplus* | *Theileria* infected *H. anatolicum* | 10 | 999 | 3.454006529 | 0.007 | 0.01 |
|  | *A. marginale* infected *R. microplus* | *Theileria* infected *R. microplus* | 9 | 999 | 4.963063677 | 0.01 | 0.01 |
|  | *Theileria* infected *H. anatolicum* | *Theileria* infected *R. microplus* | 9 | 999 | 5.322500136 | 0.004 | 0.01 |
| weighted_UniFrac | *A. marginale* infected *R. microplus* | *Theileria* infected *H. anatolicum* | 10 | 999 | 4.356141481 | 0.031 | 0.031 |
|  | *A. marginale* infected *R. microplus* | *Theileria* infected *R. microplus* | 9 | 999 | 9.578469286 | 0.017 | 0.0255 |
|  | *Theileria* infected *H. anatolicum* | *Theileria* infected *R. microplus* | 9 | 999 | 6.402764192 | 0.006 | 0.018 |

**S1; Figure 1:**
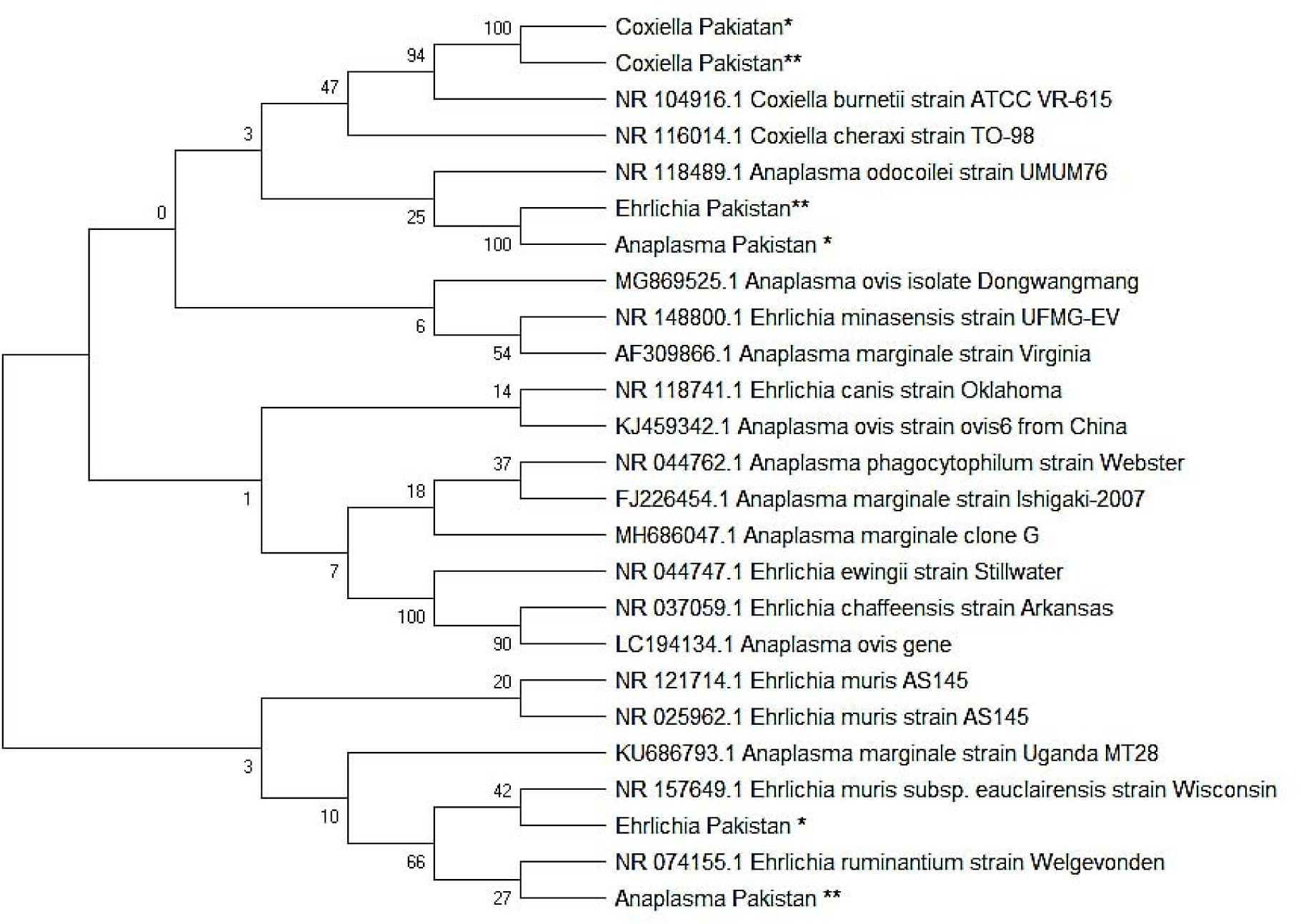
Genetic relationship between *Coxiella*, *Ehrlichia* and *Anaplasma* 16S ribosomal RNA partial sequences identified from this study against the GenBank sequences. Tree was made using the Neighbor-Joining method.

